# Isolation and characterization of cell wall and extracellular polysaccharides from cultures of the mycoparasitic strain *Tirochoderma koningiopsis*

**DOI:** 10.1101/2024.07.31.605896

**Authors:** Artur Nowak, Kamila Wlizło, Iwona Komaniecka, Monika Szymańska-Chargot, Artur Zdunek, Justyna Kapral-Piotrowska, Katarzyna Tyśkiewicz, Jolanta Jaroszuk-Ściseł

## Abstract

The *Trichoderma koningiopsis* strain showed an extracellular polymers (EPS) synthesis capacity of 1.17 g/L in an optimised Czapek-Dox medium containing sucrose (30 g/L) and yeast extract (7.5 g/L). Three fractions of wall polymers were extracted from the biomass obtained after culture: cold water soluble (WPSZ), hot water soluble (WPSC) and alkali soluble (WPSNaOH), which accounted for 13.3%, 1.8% and 20.2% of the mycelial dry weight, respectively. Structural analyses indicated that the EPS obtained was mannan, and the WPS fractions were glucans containing predominantly →4)-Glc-(1→ linked residues, with branching at →3,6)- as well as →4,6)-positions. FT-IR and FT-Raman analyses showed that α-bonds dominate in the WPSZ and WPSC fractions, whereas β-bonds dominate in the EPS and WPSNaOH fractions. The obtained polymer fractions (PS) showed antioxidant properties in the ABTS, DPPH and FRAP methods and the ability to bind bisphenol A from an aqueous environment. The most important property of the obtained PSs is their ability to reduce germination and inhibit the growth of mycelia of the phytopathogenic *Fusarium culmorum* strain. The obtained polymers exhibit a number of bioactive properties and can be used in various areas of human life.

**Highlights:**

- *Trichoderma koningiopsis* has the ability to synthesise EPS
- EPS are mainly composed of mannose and WPS of glucose
- PS have the ability to chelate BPA and have antioxidant properties
- The PS obtained has inhibitory properties against *F. culmorum*.

**Graphical Abstract:** 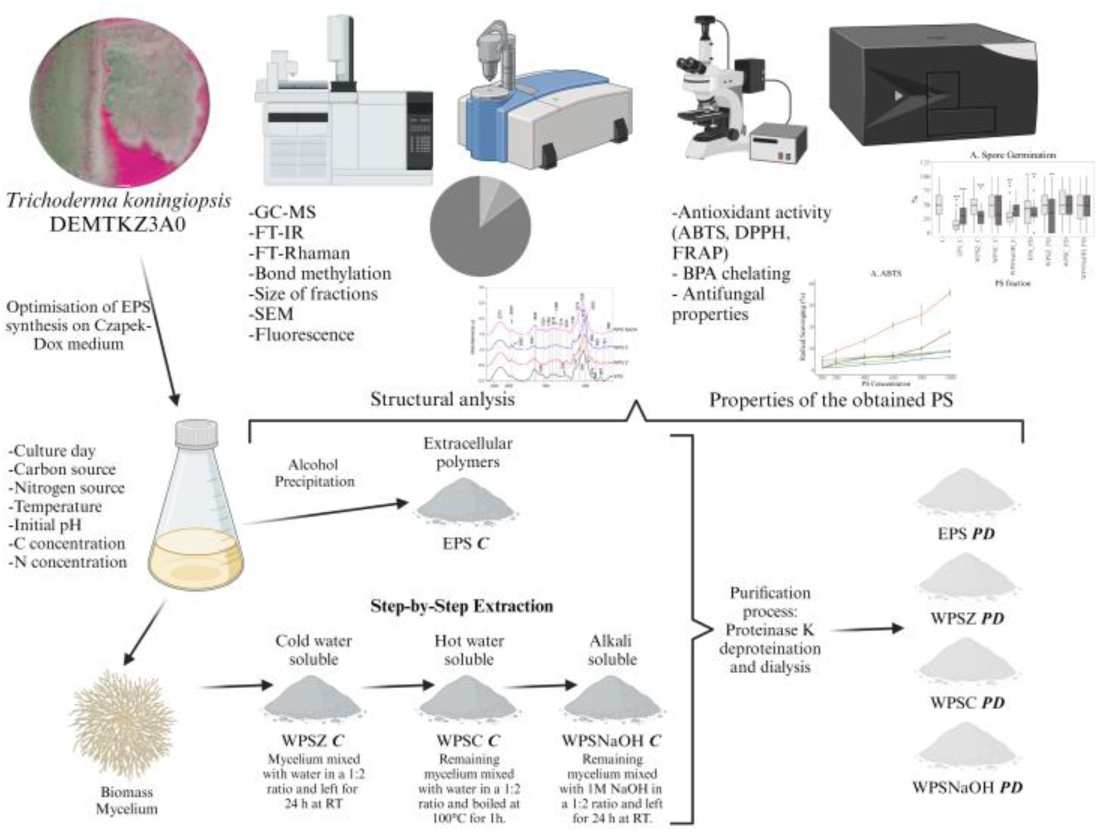

## 1. Introduction

Polymeric substances (PS) can be divided into extracellular (EPS) and cell walls (WPS) based on their location within the cell. EPS are secreted into the environment where they have a protective function against both biotic and abiotic stresses, act as a reservoir of carbon and nitrogen compounds, and have the ability to chelate ions necessary for mycelial growth and development [1]. It is known that between 40% and 95% of the composition of all extracellular polymeric substances synthesised by microorganisms are polysaccharides [2]. WPSs, on the other hand, build the cell wall and stabilise the structure of the mycelium and hyphae. They are typically composed of glucans, mannans, and chitin, interspersed with proteins, and stabilised by other substances [3]. All PS are composed of sugar monomers (e.g. glucose, mannose, galactose, and fructose) linked by α- or β-type linkages. They may consist of a single type of monomer (homopolysaccharides) or multiple monomers, known as heteropolysaccharides, and they can exhibit either a linear or branched structure [4]. In addition, proteins, phenols, amino sugars, and uronic acids may be present in the structure. The existence or non-existence of the latter component influences the electrostatic charge of polymers, resulting in either acidic or neutral properties [5]. PS can be classified according to their functions, such as sorption, surfactant, structure, and redox activity. Different combinations of these characteristics determine their properties and, thus, their roles in microbial cells [1,2]. Polymeric compounds, particularly sugar polymers, demonstrate a diverse range of bioactive properties with potential applications in medicine, environmental remediation, and agriculture [1,6–8]. In the field of medicine and pharmacy, they have been applied as compounds demonstrating anticancer, anti-inflammatory, and immunomodulatory capabilities. Notable examples of such polymers are lentinan, which is synthesised by *Lentinus edodes*, and pleuran, which is synthesised by *Pleurotus* spp. [9,10]. Furthermore, these polymers find application in the food industry as additives and in the cosmetics industry as stabilisers. Pullulan, curdlan and gellan gum are commercially available polymers exemplifying this usage [9]. Particularly intriguing are the potential applications in sustainable agriculture and environmental remediation. Microbial PSs play crucial role in stabilizing soil particles, increasing their surface area, water retention, and facilitating the formation of a matrix, which is one of the factors involved in microbe-plant interactions [11]. Moreover, the chelating properties of PS can be used to bind heavy metals or pollutants and remove them from the environment [6]. Therefore, it is important to determine the broad spectrum of properties of PSs obtained in the context of their potential human use.

In this regard, new sources of these bioactive compounds are under investigation. PS are synthesised by all groups of microorganisms, bacteria, algae, and fungi. Fungi appear to be of interest in this context due to their ability to rapidly increase biomass and produce bioactive compounds on almost any type of substrate. Polymers derived from large fruiting-body fungi, mainly belonging to Basidiomycota, have been extensively characterised and described in the literature, especially in the context of medical and pharmaceutical applications [12]. Therefore, it would be interesting to search for new sources of PS among Ascomycota strains that have already found applications in agriculture and sustainable development. Examples of such strains include species from the genus *Trichoderma*.

Fungi belonging to the genus *Trichoderma* are among the most common soil microorganisms worldwide. Various industrial and agricultural sectors make use of species within this genus as highlighted by Tyśkiewicz et al. [13]. These fungi serve as a rich source of many enzymes that are capable of degrading both cellulose polymers and chitin [14]. These include endo-xylanases, exo-glucanases, endo-glucanohydrolases, chitinases, and laccases [15]. These enzymes are classified as cell wall degrading enzymes (CWDEs). Because of their capacity to synthesise a wide range of lytic enzymes, these species are categorised as mycoparasites, which means that they live on other fungal species [16]. Another mechanism that enhances environmental adaptability and competition for ecological niches is antibiotic synthesis. A significant category of these antibiotics is referred to as peptaibols, which are polypeptide compounds with molecular weights ranging from 500 to 2200 Da. In addition, numerous Trichoderma species produce secondary metabolites belonging to terpenoids, pyrones, anthraquinones, and epipolythiodioxopiperazines [13]. In addition, *Trichoderma* spp. interact with plants to support their growth and development by synthesising phytohormones that stimulate plant growth, including indolylacetic acid (IAA), gibberellins (GA), and abscisic acid (ABA) [17]. These strains also interact endophytically with plant roots, facilitated by their ability to synthesise the enzyme deaminase, which is one of the key enzymes in controlling ET compactness in plant cells and converting it to α-ketobutyrate and ammonia [18]. All the above characteristics of the genus Trichoderma are just the tip of the potential of the species that belong to it. Consequently, they have been the focus of research as Biological Control Agents (BCAs) for many years. They are used to control fungal and bacterial diseases as well as those caused by insects and nematodes [16,19]. As a part of sustainable development and the move away from chemical pesticides, many Trichoderma species have found applications in biopreparation. Examples include *T. viride* (Anushka), *T. harzianum* CCT 7589 (Bio Zenon), *T. afroharzianum* T-22 (T-22 G Biological Fungicide), and *T. asperellum* BV-10 (Tricho-Guard) [13]. In addition to the above-mentioned properties, these strains also appear to be an interesting source of other bioactive compounds such as polymeric substances (PS). There are few reports on the ability of *Trichoderma* strains to synthesise extracellular polymers (EPS) or on the structure and properties of wall polymers (WPS) [20–22].

The aim of our work was to obtain extracellular polymers (EPS) and wall polymers (WPS) from liquid culture cultures of *Trichoderma konigiopsis* strain DEMTKZ3A0. To determine differences in structure and properties, we compared crude PS fractions (***C***) and purified fractions (***PD***). As part of our study, we optimised the culture medium, isolated EPS, and three fractions of PS (WPSZ, WPSC, and WPSNaOH), and subjected them to structural analysis. To determine the potential properties of the obtained PSs, we tested their antioxidant, chelating, and antifungal properties. The results obtained are the first report on the analysis of different PS fractions obtained from *Trichoderma konigiopsis* strain cultures.

## 2. Material and Methods

### 2.1. Trichoderma koningiopsis DEMTKZ3A0 strain

The strain *Trichoderma koningiopsis* DEMTKZ3A0 was isolated from the rhizosphere soil of winter rye (*Secale cereale* L. cv Dańkowskie Złote) grown near Lublin. The strain was morphologically and genetically determined. The sequence was subsequently deposited in the National Centre for Biotechnology Information GeneBank on 25 July 2018 under accession number MH651381 and published in the manuscript by Jaroszuk-Ściseł et.al (Jaroszuk-Ściseł et al., 2019).

### 2.2. Optimization of liquid culture

EPS synthesis yields were optimised on Czapek-Dox medium with starting composition: glucose 30 g/L, peptone 2.5 g/L, K_2_HPO_4_ 1 g/L, MgSO_4_*7H_2_O 0.5 g/L, FeSO_4_*7H_2_O, KCl 0.5 g/L, NaNO_3_ 3 g/L with initial pH 7.0 at 20°C and 120 rpm. A step-by-step optimisation was performed, which included the following:

- Culture day: 2 to 11
- Carbon sources: sucrose, glucose, fructose, mannose
- Nitrogen sources: peptone, yeast extract, NH_4_NO_3_, (NH_4_)_2_SO_4_
- Temperature: 12°C, 20°C, 28°C
- Initial pH value: 4.5; 7.0; 9.5
- Carbon source concentrations: 3.75 g/L; 15 g/L; 30 g/L;55 g/L;187.5 g/L
- Nitrogen source concentrations: 1.2 g/L; 2.5 g/L; 7.5 g/L; 20 g/L; 60 g/L

EPS was precipitated from the culture liquid using 96% ethanol 1:3 ratio at 4°C for 24h. The resulting precipitate was separated from the supernatant by centrifugation (10,000 rpm, 4°C, 14 min) and then dried on an Eppendorf Concentrator Plus vacuum evaporator (three cycles of 5 h, 30°C, 2,000 rpm, AQ mode–aqueous solutions). The resulting EPS mass was expressed in g/L.

### 2.3. Cell wall polymer isolation

After the optimization of the medium conditions, large-scale cultures were performed to obtain EPS for future studies. The mycelium obtained from the culture was used as a source for cell wall polymers extraction, according to the methods of Palacios et al. with some modifications [24].

Following the separation from the culture liquid, the obtained mycelium was rinsed four times with dH_2_O to remove ballast substances, and subsequently underwent lyophilization. The dried mycelia were ground in a blade mill to obtain a homogeneous powder. To remove low molecular weight substances, mycelium was washed four times with methanol (MeOH) and then dried. Three fractions of cell wall polymers were extracted from the purified mycelia using a step-by-step method:

1. Cold water soluble: WPSZ – The resulting mycelium was mixed with water in a 1:2 ratio and left for 24 h at RT.
2. Hot water soluble: WPSC – The remaining mycelium fraction obtained in step 1 was mixed with water in a 1:2 ratio and boiled at 100°C for 1h.
3. Alkali soluble: WPSNaOH – The remaining mycelium fraction obtained in step 2 was mixed with 1M NaOH in a 1:2 ratio and left for 24 h at RT.

All extraction steps were performed 2 times. Between each step, the mycelia was lyophilised and the percentage of each fraction was determined. After separation of the supernatant, the polymers were precipitated using ethanol at a ratio of 1:3. In the case of the WPSNaOH fraction, the supernatant was neutralised with 1M HCl to pH ∼7.0 before precipitation. All WPS fractions obtained were lyophilised.

### 2.4. Purification of obtained polymers

All the obtained Polymer (PS) fractions (EPS and WPS) were subjected to deproteinization using proteinase K. Samples of the obtained polymers were suspended in phosphate buffer pH 7.5 with 0.01% NaN_3_. The reaction mixture consisted: 10 mL of buffer per 1 g of PS fraction and 1 mg of proteinase K. Deproteinization was carried out with continuous stirring at 37°C for 72 h. Afterwards, the suspensions were heat-inactivated at 80°C for 15 min. The obtained preparations were dialysed against distilled water on a cellulose membrane with a pore size of 12 kDa for 7 days. Subsequently, all the polymer fractions were lyophilised [25].

In this manuscript, the crude fraction of polymers is described as ***C***, and the purified fraction is referred to as ***PD***.

### 2.5. Compositional analysis of polymers

#### 2.5.1. Sugar composition of EPS and WPS fractions

Polymers fractions were hydrolysed in 2M TFA for 4 h at 100°C, dried, and converted in (amino)alditol acetates, by performing N-acetylation, reduction using sodium borodeuteride, and acetylation procedure [26]. The samples purified by a chloroform : water (1:1, v/v) extraction. Peracetylated (amino)alditol acetates were recovered from the organic phase and dried using an anhydrous sodium sulphate. Samples were analysed by gas chromatography coupled to a mass spectrometer (GC-MS) technique. Twenty micrograms of inositol was added to each sample before hydrolysis, as an internal standard (concentration 1.026 µg/µL).

#### 2.5.2. Methylation analysis of EPS and WPS fractions

The linkage position between sugar components in polymers preparations (purified fractions, ***PD***) was established by performing methylation analysis using iodomethane as a methylation agent according to the Ciucanu and Kerek procedure [27]. Methylated products were extracted into the chloroform, acid hydrolyzed (2 M trifluoroacetic acid, 100°C, 4 h), reduced with sodium borodeuteride, and acetylated [26]. Samples were analysed using GC-MS technique.

#### 2.5.3. Gas chromatography-mass spectrometry (GC-MS) analysis of sugar derivatives

Obtained amino-alditol acetates, and partly methylated alditol acetates were analysed by GC-MS using a gas chromatograph (7890A, Agilent Technologies, Inc., Wilmington, DE, USA) connected to a mass selective detector (MSD 5975C, inert XL EI/CI, Agilent Technologies, Inc., Wilmington, DE, USA). The chromatograph was equipped with an HP-5MS column (30 m x 0.25 mm). Carrier gas was helium, with a flow rate of 1 ml min^-1^. The temperature program was as follows: 150°C for 5 min, increased to 310°C with a rise of 5°C min^−1^; and the final temperature was kept for 10 min. The components were identified based on their characteristic mass spectra and retention time, and compared with known analytical standards.

#### 2.5.4. Gel permeation chromatography (GPC) of EPS and WPS fractions

The average molecular mass of polysaccharides in PS fractions were determined using gel permeation chromatography (GPC) with a Sepharose CL 6B column (0.7 cm x 90 cm). The polysaccharide preparations (5 mg) were dissolved in 1M sodium hydroxide solution and eluted from the column with a 1M NaOH, with a flow rate of 0.2 ml/min. Fractions containing 40 droplets (∼1 ml) were collected using a BioRad model 2110 collector into a dedicated 6 ml PP tubes [28].

Carbohydrate content in all fractions was determined using Dubois method [29] and dextran of known molecular masses were used as standards to calibrate the column.

#### 2.5.5. FT-IR and Raman spectroscopy of EPS and WPS fractions

FT-IR spectra were collected on a Nicolet 6700 FT-IR (Thermo Scientific, Madison, WI, USA). The Smart iTR ATR sampling accessory was used. The spectra was collected in the range of 4000–650 cm^−1^. For each material, three samples under the same conditions were examined. For each sample, 200 scans were averaged with a spectral resolution of 4 cm^−1^. Then for a given material, a final average spectrum was calculated and baseline correction was performed. These spectra were normalized to 1.0 at band ca. 1020 cm^-1^.

The FT-Raman spectra were acquired on an FT-Raman module (NXR FT Raman) for a Nicolet 6700 Fourier transform infrared spectroscopy bench using a InGaAs detector and CaF_2_ beam splitter (Thermo Scientific, Madison, WI, USA). The samples in the form of powder were placed in stainless cubes and were illuminated using Nd:YAG excitation laser operating at 1064 nm. The maximum laser power was 0.6W. The spectra were recorded over the range of 3500–250 cm^-1^, and each spectrum has an average of 200 scans with 8 cm^-1^ resolution. The analyzed spectra were averaged over three registered spectra. Baseline correction and normalization to 1.0 at band ca. 2930 cm^-1^ was used.

All the spectra manipulation was performed in OMNIC Software (Thermo Scientific), while the graphical presentation was obtained in Origin Software (Origin Lab v8.5 Pro, Northampton, USA).

#### 2.5.6. Microscopic visualization

##### Fluorescence microscopy

Calcofluor (200 µL) was added to 5 mg of test PS fractions and incubated for 25 min in the dark. The solution was then centrifuged (10 min at RT at 10,000 rpm) and the resulting sediment was transferred to a glass slide and observed using an Olympus BX53 Microscope equipped with an Olympus XC30 camera using excitation at 365 nm and emission at 435 nm.

##### Scanning electron microscopy (SEM)

The dried samples were mounted on metal stubs with a piece of double-coated carbon conductive tape and coated with gold using an Emitech K550X sputter coater. Scanning electron micrographs were obtained using a scanning electron microscope (TESCAN Vega 3 LMU microscope, Czech Republic). Scanning electron microscopy (SEM) images were captured at a magnification of 1000× based on a voltage of 30kV acceleration under high vacuum conditions.

### 2.6. Properties of the polymers obtained

For all antioxidant assays, PS fractions were prepared in the concentration range of 100-1000 ug/mL in terms of soluble sugars. Sugar concentration was determined by the Dubois method [29].

#### 2.6.1. Antioxidant properties

##### 2,2-azinobis (3-ethyl-benzothiazoline-6-sulfonic acid) – ABTS

The stock solution was prepared 12 h before the assay by dissolving 76 mg ABTS in 20 mL PBS buffer and adding 13.2 mg K_2_S_2_O_8_. The absorbance of the resulting solution was set to ABS_734_=0.7. Ten microlitres of the sample was then mixed with 990 µL of solution and incubated for 5 min at RT. The absorbance at 734 nm was measured, and the ABTS activity was calculated using the following formula:

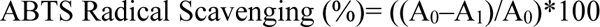

where: **A_1_** – absorbance of the sample, **A_0_** – absorbance of the blank [30].

##### 2,2-diphenyl-1-picrylhydrazyl – DPPH

A working solution was prepared immediately before measurement by dissolving DPPH in 96% ethanol at a concentration of 0.2 mg/mL. Then, 100 µL of the test sample was mixed with 100 µL of the reagent and incubated for 10 min at RT. The absorbance at 515 nm was measured, and the DPPH activity was calculated using the following formula:

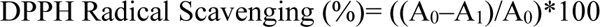

where: **A_1_** – absorbance of the sample, **A_0_** – absorbance of the blank [30].

##### Ferric Reducing Antioxidant Power – FRAP

Immediately before the assays, FRAP reagent was prepared containing 300 mM acetic buffer (pH 3.6), 10 mM TPTZ (2,4,6-tripyridyl-s-triazine) in 40 mM hydrochloric acid, and 20 mM FeCl_3_ in a 10:1:1 ratio. Twenty microlitres of the sample were mixed with 150 µL of reagent and incubated for 10 min at RT. A calibration curve (ascorbic acid) and reagent sample (blank) were prepared in parallel. The absorbance at 593 nm was measured, and the FRAP activity was calculated according to the following formula:

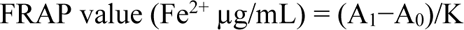

where: **A_1_** – absorbance of the sample, **A_0_** – absorbance of the blank, **K** – coefficient of the calibration curve (K=0.013) [31].

#### 2.6.2. Chelating capacity – BPA

Samples of 10 mg of each PS fractions were placed into 2 mL Eppendorf tubes, mixed with 0.5 mL of dH_2_O, and left overnight at temperature of 25°C. Afterwards, 0.5 mL of 160 mg/L bisphenol A (BPA) water solution was mixed with each sample. The mixture was shaken at temperature of 25°C and 800 rpm for 30 minutes using a JWE Electronics WL-972S (Poland). After incubation, the samples diluted in water were mixed with 1 mL of 96% ethanol, to precipitate them, apart from samples of WPS NaOH ***C*** and WPS NaOH ***PD***, to which 1 mL of deionised water was added. All samples were then centrifuged for 10 minutes at 13.400 RPM, and the supernatant was transferred to a 96-well UV-permeable plate. Starting solution of 80 mg/L BPA diluted two times in 96% ethanol (control for water-diluted fractions) and diluted two times in water (control for alkaline fractions) was also transferred, to evaluate its maximum absorption. Assessed and control samples of supernatants and BPA solutions were measured on a microplate reader (Multiskan Go 1510 Microplate Reader, Thermo Fisher Scientific, USA) at 275 nm. To calculate the rate of BPA removal was using the equation:

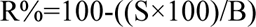

where: **R%** – removal %, **S** – supernatant absorbance, **B** – BPA starter solution

#### 2.6.3. Inhibition of *Fusarium culmorum* spore germination and hyphe lenght

Suspension of *F. culmorum* DEMFc37 phytopathogenic fungi spores was prepared in CFU=2.0*10^6^/mL concentration. Subsequently, 250 µL of the spores and 250 µL of the aqueous PS fractions were transferred to Eppendorf tubes to obtain final PS concentrations of 0.05% and 0.1%, respectively. The samples were incubated for six days at 25°C with shaking at 120 rpm. Then, 30 µL of the suspension was transferred to a glass slide and observed using an Olympus BX53 Upright Microscope equipped with an Olympus XC30 camera. The examination involved observations at 45 randomly selected points on each slide. The number of spores germinated (%) and the average length of the hyphae (µm) was determined using ImageJ software [32].

*F. culmorum* DEMFc37 was isolated in Poland near Lublin region from winter wheat (*Triticum aestivum* L.) plants with severe fusariosis symptoms. Strains were deposited at in the Centraalbureau voor Schimmelcultures Collection (CBS). The *F. culmorum s*train was morphologically and genetically identified according ITS region sequencing, which were deposited in the National Centre for Biotechnology Information GeneBank at 10 April 2006 with a DQ450878 sequence number [33].

### 2.7. Chemicals

All reagents used in the assays were purchased from: POCH, Gliwice, Poland; Sigma-Aldrich, Hamburg, Germany; Chempur, Piekary Śląskie, Poland, BTL, Łódź, Poland and Merck, Darmstadt, Germany.

### 2.8. Statistical analysis

All the samples were separated into three independent replicates, which were then assayed in three independent assays, yielding a replicate count of n=3. Data are presented as mean values with standard deviation (SD). The results were subjected to analysis of variance (ANOVA) followed by Tukey’s post hoc test for multiple comparisons at p < 0.05 and t-Student two sample tests in some analysis. The data were also analysed using PCA (Principal Component Analysis). All tests were performed using the open-source software RStudio for Windows version 2023.03.0 + 386 (Posit, PBC, GNU Affero General Public Licence v3).

## 3. Results

### 3.1. Isolation polymers from *T. koningiopsis* strain cultures

The *T. koningiopsis* strain was able to synthesise EPS in liquid cultures. A modified Czapek-Dox medium with a final sucrose concentration of 30 g/L and yeast extract concentration of 7.5 g/L proved to be optimal. The maximum EPS synthesis was reached on the third day of the culture. Temperature and initial pH values did not affect the amount of EPS obtained (Table 1). During optimisation, an increase in EPS synthesis of 40% from 0.78 g/L to 1.17 g/L was achieved. The test strain was a fast-growing species, and the average biomass gain under optimal conditions was 13 g/L. Despite inhibition of EPS synthesis in some culture variants, mycelial growth was not inhibited (Table S1).

**Table 1.** Results of step-by-step optimisation of *T. koningiopsis* strain culture on the amount of EPS obtained (g/L). Statistical analysis was performed by Anova test using Tukey post hoc p<0.05.

| EPS<br>yield<br>g/L | Optimised parameter |  |  |  |  |  |  |  |  |  |
| --- | --- | --- | --- | --- | --- | --- | --- | --- | --- | --- |
|  | Day of culture |  |  |  |  |  |  |  |  |  |
|  | 2 | 3 | 4 | 5 | 6 | 7 | 8 | 9 | 10 | 11 |
|  | 0.55±0.07<br>ab | **0.82±0.11**<br>a | 0.4±0.06<br>bc | 0.04±0.04<br>e | 0.08±0.05<br>de | 0.2±0.14<br>cde | 0.27±0.08<br>cd | 0.16±0.07<br>de | 0.22±0.1<br>cde | 0.13±0.05<br>de |
|  | Carbon source |  |  |  |  |  |  |  |  |  |
|  | Sucrose |  |  | Glucose |  |  | Fructose |  | Mannose |  |
|  | **0.89±0.11**<br>a |  |  | 0.7±0.12<br>b |  |  | 0.24±0.03<br>c |  | 0.7±0.1<br>b |  |
|  | Nitrogen source |  |  |  |  |  |  |  |  |  |
|  | Peptone |  |  | Yeast extract |  |  | NH<sub>4</sub>NO<sub>3</sub> |  | (NH<sub>4</sub>)<sub>2</sub>SO<sub>4</sub> |  |
|  | 0.9±0.1<br>b |  |  | **1.09±0.12**<br>a |  |  | 0.79±0.04<br>bc |  | 0.66±0.02<br>c |  |
|  | Temperature (°C) |  |  |  |  |  |  |  |  |  |
|  | 12 |  |  | 20 |  |  | 28 |  |  |  |
|  | 0±0<br>b |  |  | **0.95±0.13**<br>a |  |  | 0±0<br>b |  |  |  |
|  | Initial pH value |  |  |  |  |  |  |  |  |  |
|  | 4.5 |  |  | 7.0 |  |  | 9.5 |  |  |  |
|  | 1.07±0.08<br>a |  |  | **1.05±0.02**<br>a |  |  | 0±0<br>b |  |  |  |
|  | Carbon source concentrations (g/L) |  |  |  |  |  |  |  |  |  |
|  | 3.75 |  | 15 |  | 30 |  | 55 |  | 187.5 |  |
|  | 0.42±0.02<br>bc |  | 0.53±0.03<br>b |  | **1.01±0.1**<br>a |  | 0.33±0.01<br>c |  | 0.37±0.02<br>c |  |
|  | Nitrogen source concentrations (g/L) |  |  |  |  |  |  |  |  |  |
|  | 1.2 |  | 2.5 |  | 7.5 |  | 20 |  | 60 |  |
|  | 0.12±0.01<br>d |  | 0.65±0.09<br>b |  | **1.17±0.11**<br>a |  | 0.66±0.06<br>b |  | 0.21±0.04<br>c |  |

The cell wall of the *T. koningiopsis* strain consisted of 13.3% cold-soluble polymers (WPSZ), 1.8% hot-soluble polymers (WPSC) and 20.2% alkali-soluble polymers (WPSNaOH). The remaining precipitate consisted of insoluble polymers (64.7%) (Fig. 1).

**Fig. 1.**
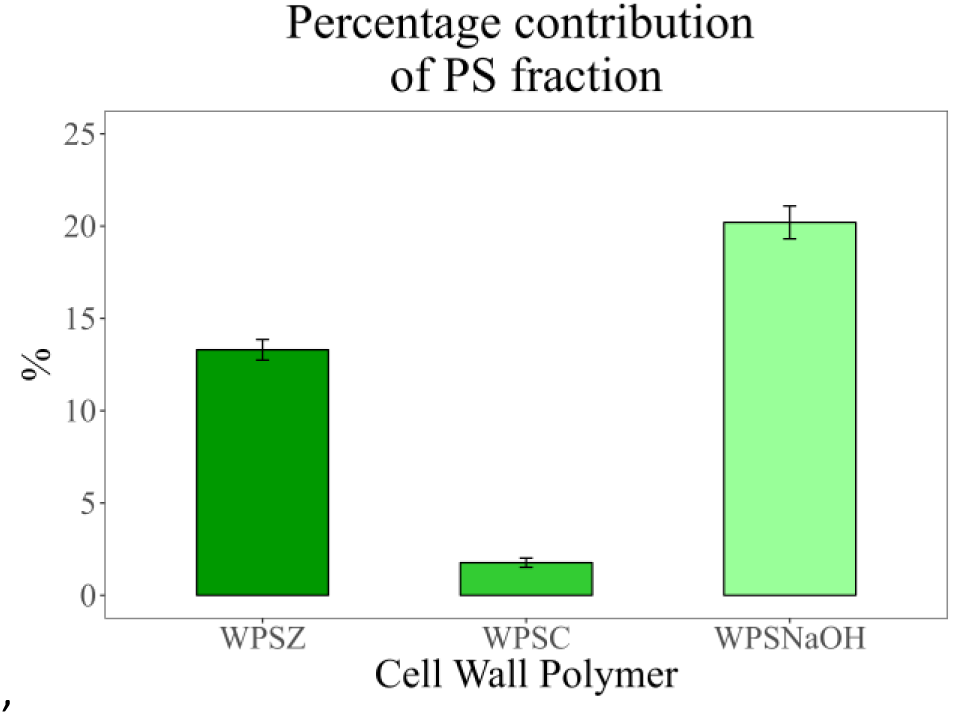
Percentage of the cell wall polymer fractions in the biomass of *T. koningiopsis* cuttings

### 3.2. Compositional analysis of obtained polymers

Sugar analysis of the obtained PS fractions revealed that all the cell wall polymers had a predominance of glucose (Glc), accounting for approximately 60 % to 85 % of the total sugar components (Fig. 2). In the case of the WPSC (Fig. 2E,F) and WPSNaOH (Fig. 2G,H) fractions, a considerable increase in the percentage of Glc was observed in the samples after purification. It is interesting to note that the predominant monomer in the crude EPS (***C***) fraction was glucose (72.3%) and after deproteinization and dialysis (***PD*** preparation) the main component was mannose (79.2%) (Fig. 2A,B). In crude EPS two hexosamines (GlcN and GalN) were identified, but in ***PD*** preparation they were not noticed. Moreover, in the WPSZ ***C*** fraction, a small amounts of pentoses identified as ribose (1.5 %) and xylose (0.4 %), were also detected.

**Fig. 2.**
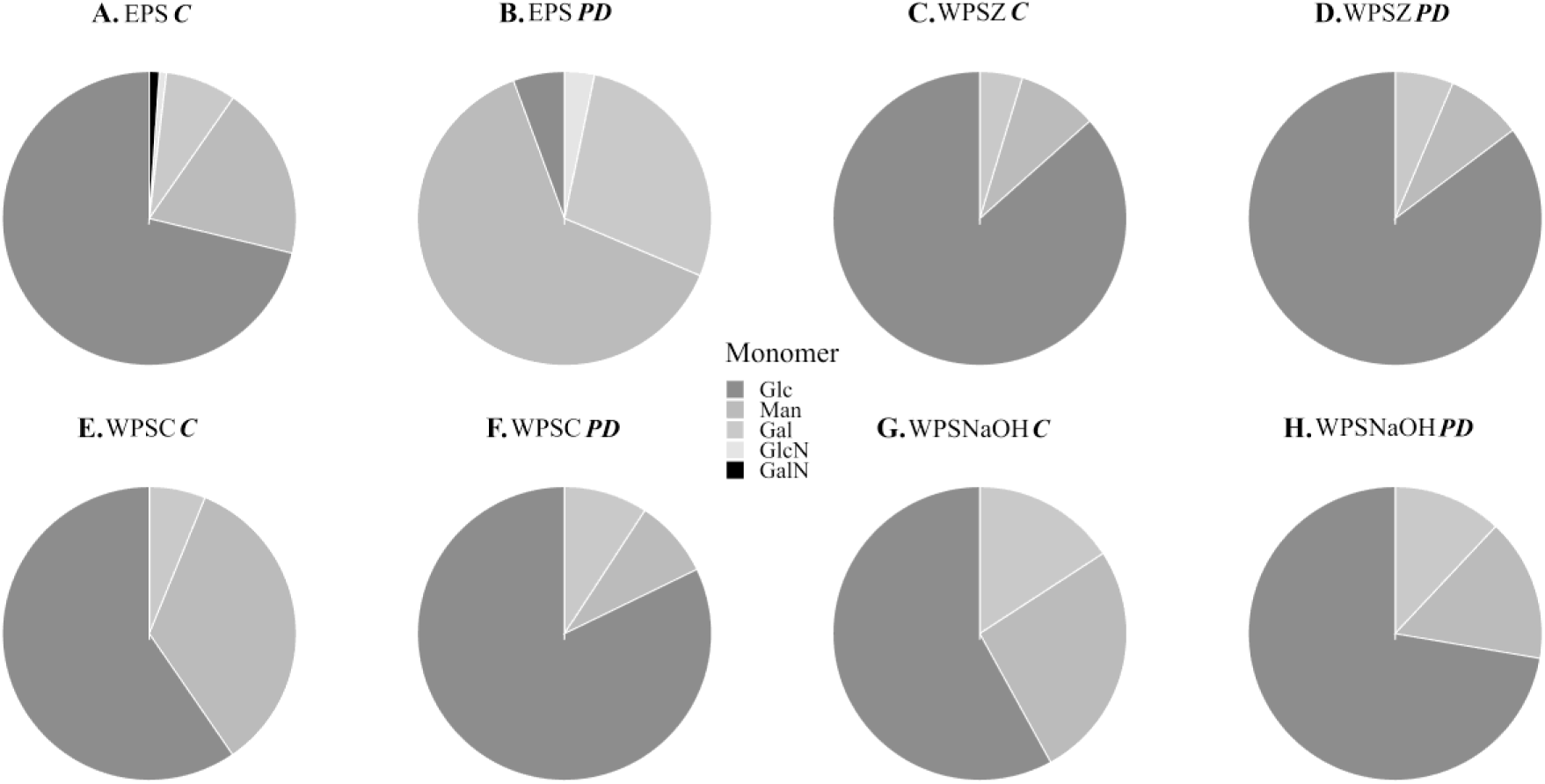
Analysis of the composition of sugar monomers in the polymer fractions obtained from *T. koningiopsis* performed before (***C***) and after (***PD***) the purification process. Glc – Glucose, Man – Mannose, Gal – Galactose, GlcN – Glucosamine, GalN – Galactosamine.

Linkage analysis was conducted on all ***PD*** preparations. Within EPS ***PD,*** only four forms of methylated sugar derivatives were present: terminal Man, →4)-linked form of hexose II, and two different branched →3,6)-linked hexoses. In WPSZ all three hexoses (Man, Glc and Gal) were present as terminal residues, but →4)-linked hexose II (most probably Glc) strongly predominated, constituting 77.6 % of the preparation). Additionally, →4,6)- and →3,6)-branched hexose forms were also present. Methylation analysis of WPSC showed the presence of two terminal hexoses (Glc and Man), also →3)-linked, two →4)-linked, →2)- linked and →6)-linked hexoses. Moreover branched forms were also present, namely: →3,4)- and two →3,6)- linked hexoses. In WPSNaOH all three hexoses (Man, Glc and Gal) were present as terminal residues, but →4)- linked hexose (Glc) strongly predominated. Additionally, →4,6)- and two →3,6)-branched hexose forms were also present in this sample (Table 2).

**Table 2.**
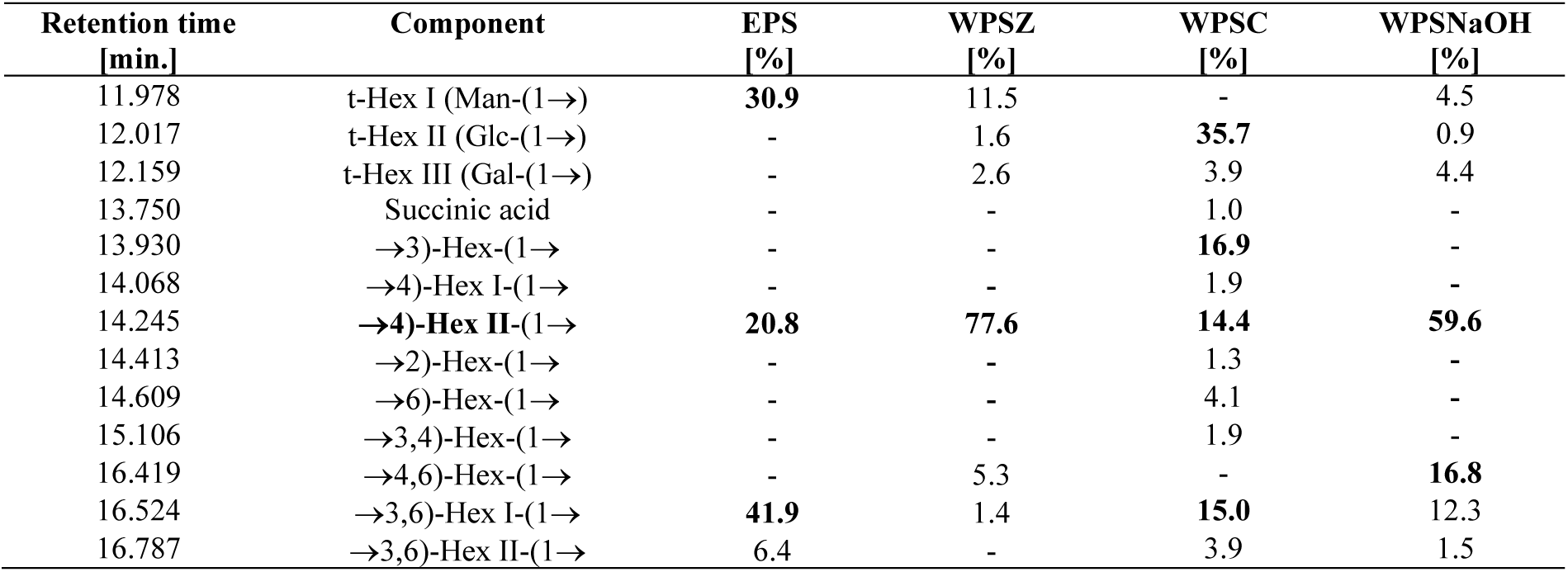
Linkage analysis of PS preparations (PD) obtained from *T. koningiopsis*. PS samples were methylated, hydrolyzed, reduced and acetylated. The obtained partly methylated alditol acetates were identified by GC–MS, based on their characteristic mass spectra and retention times. Predominant components were marked in bold.

| Retention time<br>[min.] | Component | EPS<br>[%] | WPSZ<br>[%] | WPSC<br>[%] | WPSNaOH<br>[%] |
| --- | --- | --- | --- | --- | --- |
| 11.978 | t-Hex I (Man-(1→)) | **30.9** | 11.5 | - | 4.5 |
| 12.017 | t-Hex II (Glc-(1→)) | - | 1.6 | **35.7** | 0.9 |
| 12.159 | t-Hex III (Gal-(1→)) | - | 2.6 | 3.9 | 4.4 |
| 13.750 | Succinic acid | - | - | 1.0 | - |
| 13.930 | $\rightarrow 3$)-Hex-(1→)) | - | - | **16.9** | - |
| 14.068 | $\rightarrow 4$)-Hex I-(1→)) | - | - | 1.9 | - |
| 14.245 | **$\rightarrow 4$)-Hex II-(1→))** | **20.8** | **77.6** | **14.4** | **59.6** |
| 14.413 | $\rightarrow 2$)-Hex-(1→)) | - | - | 1.3 | - |
| 14.609 | $\rightarrow 6$)-Hex-(1→)) | - | - | 4.1 | - |
| 15.106 | $\rightarrow 3,4$)-Hex-(1→)) | - | - | 1.9 | - |
| 16.419 | $\rightarrow 4,6$)-Hex-(1→)) | - | 5.3 | - | **16.8** |
| 16.524 | $\rightarrow 3,6$)-Hex I-(1→)) | **41.9** | 1.4 | **15.0** | 12.3 |
| 16.787 | $\rightarrow 3,6$)-Hex II-(1→)) | 6.4 | - | 3.9 | 1.5 |

All FT-IR spectra of EPS and WPS from fungi contain bands at ca. 3273 cm^-1^ and in the range 3000-2800 cm^-1^ and are assigned to OH and CH groups vibrations. The ***PD*** treatment didn’t influence those bands. Crude (***C***) sample spectra contained intense bands characteristic of proteins. In the case of FT-IR spectra it was 1647 cm^-1^ stretching vibration of C=O and C-N in Amid I, 1550 cm^-1^ stretching vibration of C-N and bending vibration of N-H in Amid II (both related to peptide bands CO-NH), and 1340-1240 cm^-1^ vibration of C-N in Amid III (Fig. 3A,B). While for FT-Raman spectra the bands characteristic for β turn at 1670 cm^-1^ and for β sheet at 1628 cm^-1^ in Amid I were detected, while in the range 1300-1230 cm^-1^ bands characteristic for α helix, β sheet, and random coil in Amid III can be found (Fig. 3C). In FT-Raman spectra of all WPS fractions the band at 1003 cm^-1^ and characteristic for phenylalanine was detected. While for EPS spectrum the band at 992 cm^-1^ and characteristic for phenylalanine ring vibration was present (Fig. 3C). All abovementioned bands diminished or even disappeared for ***PD*** samples spectra (Fig. 3D). It is important to note that both FT-IR and FT-Raman spectra below 1700 cm^-1^ contain bands characteristic for carbohydrates.

**Fig. 3.**
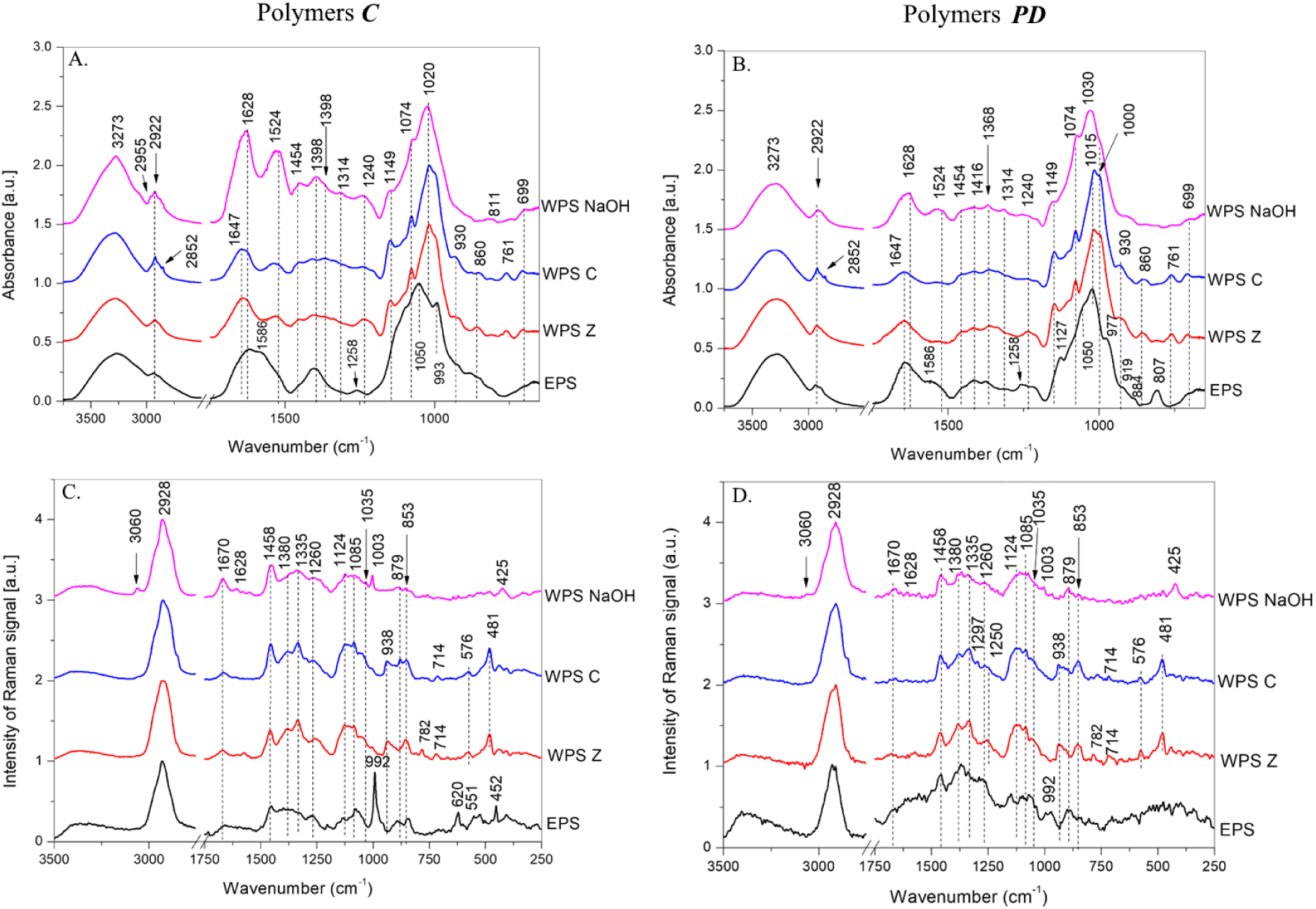
FTIR (A,B) and FTRaman (C,D) spectra of fractions isolated from *T. koningiopsis,* studied before (C) and after (PD) the purification process.

The average molecular mass of polysaccharides in ***PD*** fractions was determined using GPC chromatography. The highest molecular weight fractions were observed for WPSC and WPSNaOH, corresponding to range 831.76 - 158.49 kDa and 831.76 - 691.83 kDa, respectively. A low-molecular-weight fraction was also present in both preparations. In contrast, the WPSZ consisted of only one fraction with a range of 575.44 - 398.11 kDa. The smallest polymers were observed in the case of EPS, where separation yielded three fractions with masses of 52.48, 25.12 and 2.75 kDa, respectively (Table 3).

**Table 3.** Fraction size of preparations (*PD*) isolated from *T. koningiopsis*, established using GPC at Sepharose CL 6B column using 1 M sodium hydroxide as an eluent.

| Polymer | Polymer size [kDa] |  |  |
| --- | --- | --- | --- |
|  | High molecular weight<br>(1 000 – 100 kDa) | Medium molecular weight<br>(100 – 10 kDa) | Low molecular weight<br>(<10 kDa) |
| **EPS** |  | 52.48; 25.12 | 2.75 |
| **WPSZ** | 575.44 – 398.11 |  |  |
| **WPSC** | 831.76 – 158.49 |  | 2.29 |
| **WPSNaOH** | 831.76 – 691.83 | 14.45 |  |

Upon visualisation of the resulting preparations, the presence of polymers containing cellulose-like or chitin-like bonds capable of interacting with calcofluor was observed. An increase in the fluorescence intensity was observed for all purified polymers (PS) (Fig. 4).

**Fig. 4.**
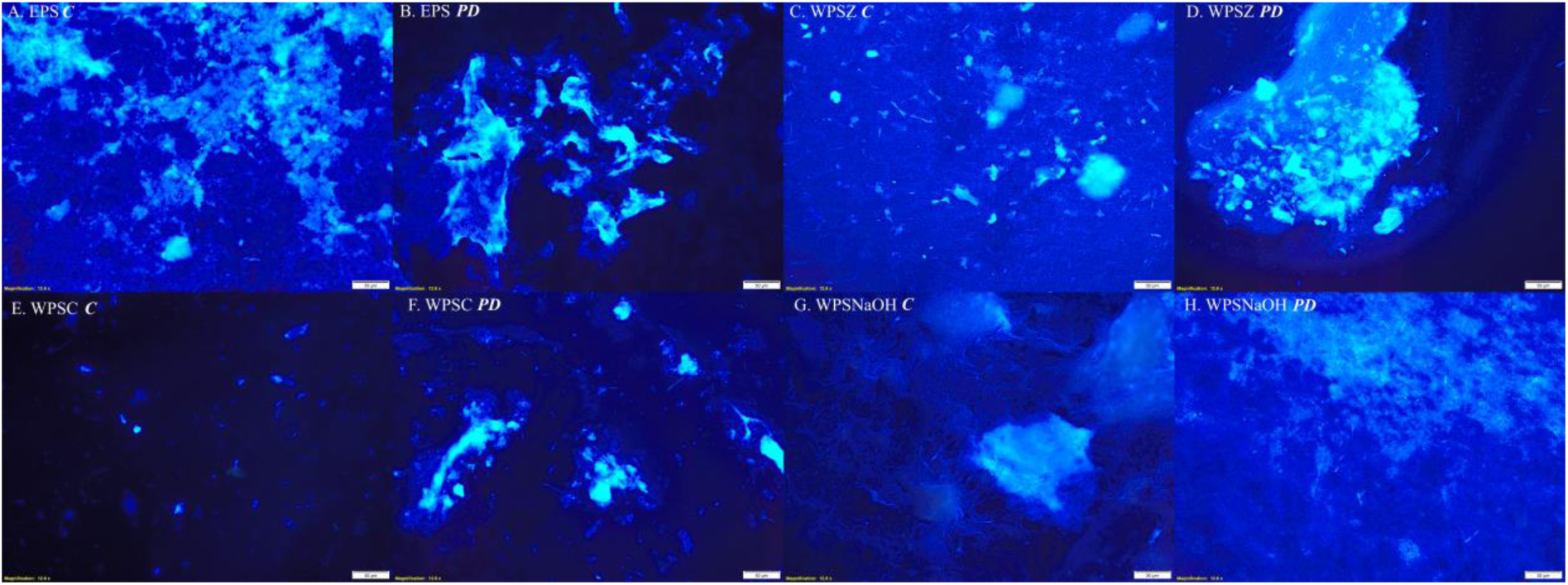
Visualisation of sugar compounds in the polymer fractions studied before (***C***) and after (***PD***) the purification process.

During the SEM analysis, it was noted that the process undergone by the polymers prior to purification was more homogeneous and compact, forming irregular shapes resembling crystals. The purification process, on the other hand, increased the porosity of the PS studied and the formation of network-like structures (Fig. 5).

**Fig. 5.**
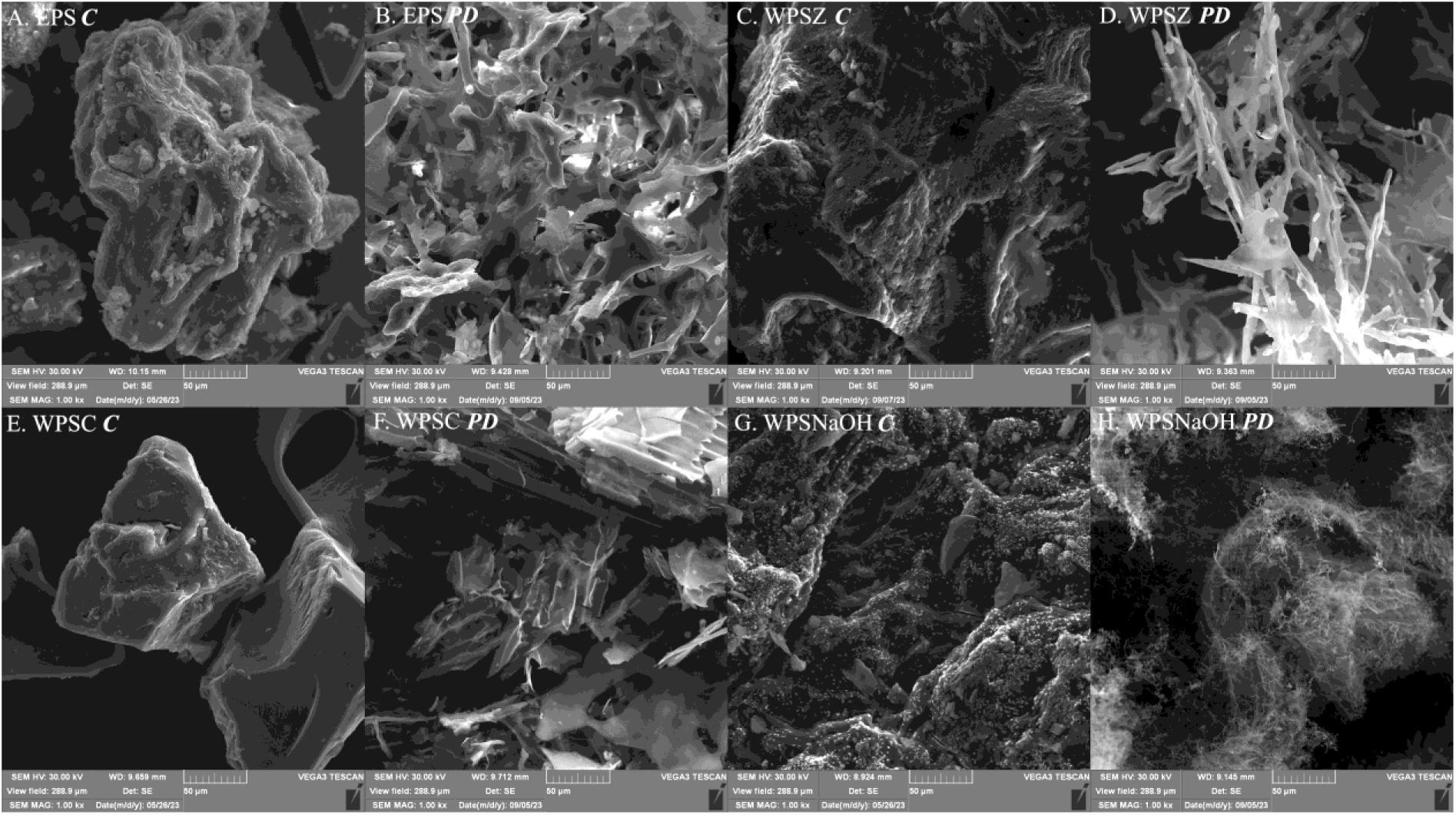
SEM images of polymer fractions studied before (***C***) and after (***PD***) the purification process at x1000 magnification.

### 3.3. Properties of obtained polymers

All examined polymers tested showed antioxidant properties, which varied depending on the methodology employed and the PS fraction (***C*** or ***PD***). In the ABTS assay, WPSNaOH ***C*** showed the greatest antioxidant properties at a concentration of 1000 µg/mL, with a free radical scavenging level of 35% (Fig. 6A). However, in the DPPH assay, a similar level of free radical scavenging was achieved by the EPS PD fraction. In this method, an increase in activity was observed in the PS fractions subjected to purification (***PD***) (Fig. 6B). In contrast, the water-soluble wall polymer fractions WPSZ ***PD*** and WPSC ***PD*** showed the highest ability to reduce Fe^3+^ ions at concentrations of 4 and 6 µg/mL, respectively (Fig. 6C).

**Fig. 6.**
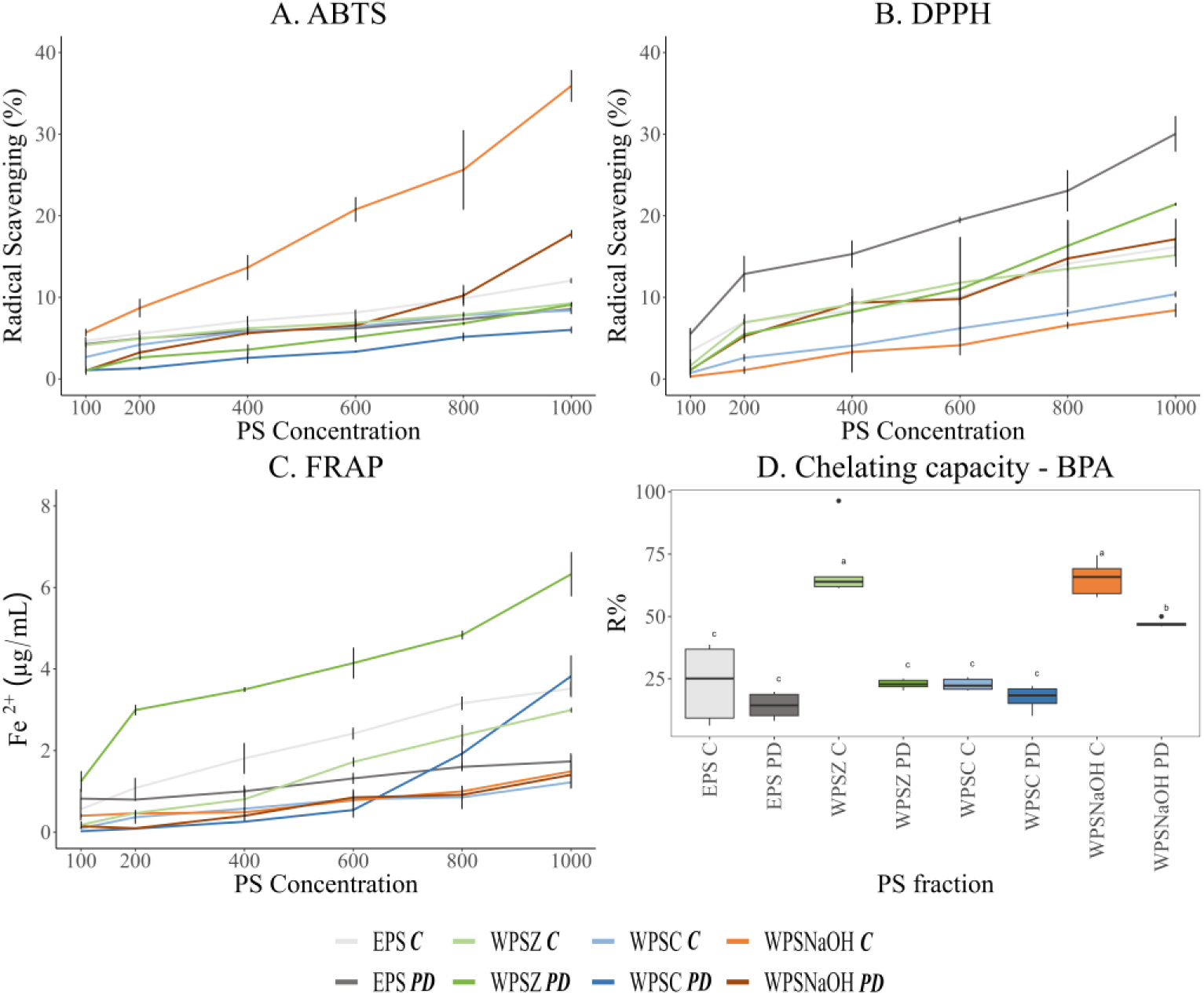
Antioxidant properties of the PS obtained. A. ABTS, B. DPPH, C. FRAP and D. BPA binding capacity

The obtained polymers showed bisphenol A binding capacities ranging from 25 to 70%. However, the highest chelating capacity was demonstrated by the friction of WPSZ ***C***, WPSNaOH ***C***, and WPSNaOH ***PD*** at levels of 68%, 65%, and 47%, respectively (Fig. 6D).

The EPS fractions (both EPS ***C*** and EPS ***PD***) had the greatest effect on spore germination of the *F. culmorum* strain. In the case of EPS ***C***, a statistically significant (p < 0.001) decrease in the number of germinated spores was observed at both the 0.05% and 0.1% concentrations. The EPS ***PD*** fraction also significantly reduced the number of germinated spores; however, a greater reduction was observed at 0.1% (p<0.01) than at 0.05% (p<0.05) (Fig. 7A). The inhibition of spore germination did not correlate with a reduction in the length of the outgrown hypea of the phytopathogenic strain. Significant effects on the length of the hypea were observed for the WPSC ***C*** and WPSNaOH ***C*** fractions at both concentrations, and for WPSZ ***PD*** at 0.1% (Fig. 7B).

**Fig. 7.**
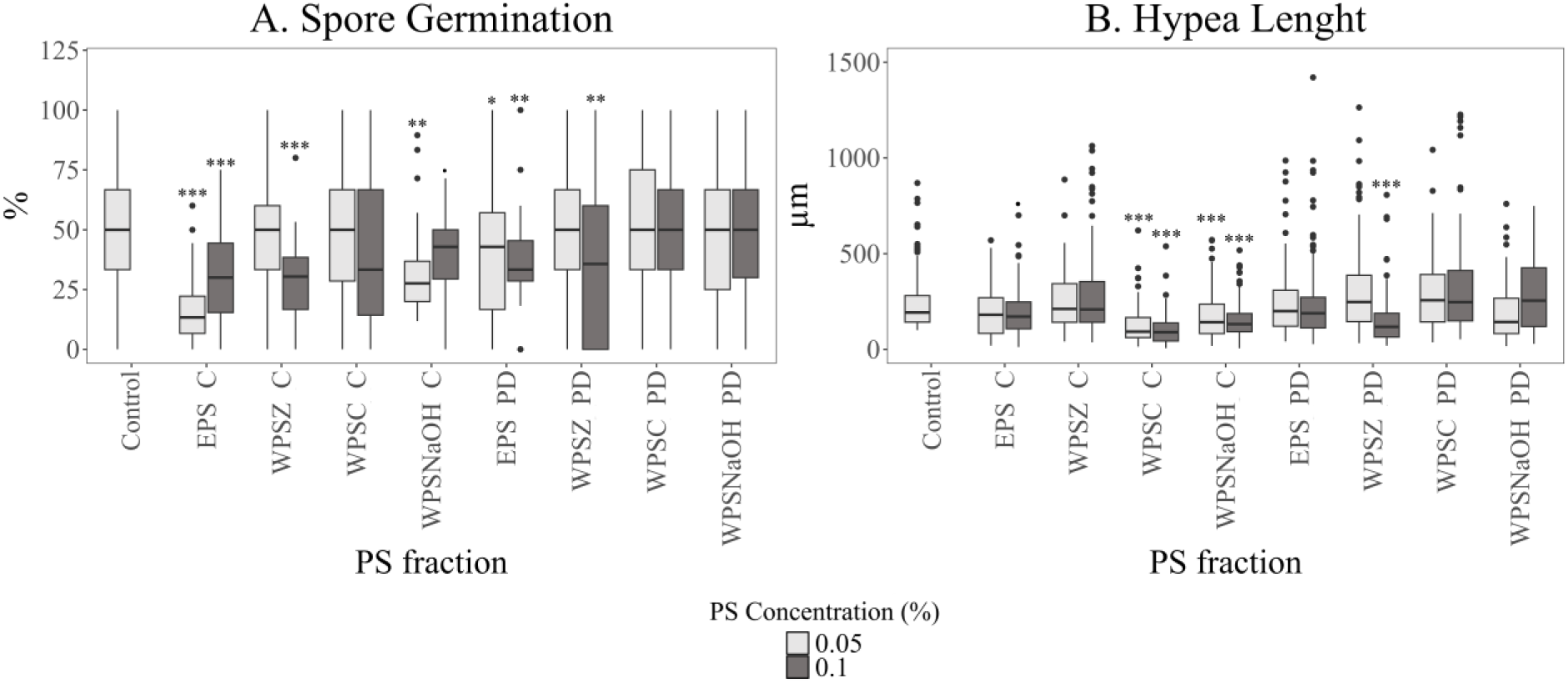
Effect on A. Spore germination; B. Stem length of *F. culmorum* strain Fc37 after a six-day incubation in the presence of 0.05% and 0.1% suspensions of the PS tested. Significance against control determined by Dunnett’s test, where p value is: **\*\*\*** 0.001; **\*\*** 0.01; **\*** 0.05;. 0.1.

### 3.4. Statistical analysis

The correlation between the amount of EPS and various optimisation parameters was determined by Principal Component Analysis (PCA). The diagram shows the influence of the culture day (day), carbon (C) and nitrogen (N) source, temperature (T), pH (pH) and concentration of the carbon (Ccon) and nitrogen (Ncon) source on the biomass and the efficiency of EPS synthesis by the *T. koningiopsis* strain (Fig. 8A). The first two dimension of PCA explained 100% of the total variation, with principal component 1 (Dim1) accounting for 48.8% and principal component 2 (Dim2) accounting for 51.2% of the variance. EPS synthesis by the tested strain was most influenced by the type of C and N sources, their concentration, and the initial pH value of the medium. The highest EPS synthesis was observed with sucrose (C_suc), glucose (C_glc), and mannose (C_mann) as carbon sources and peptone (N_pep) and yeast extract (N_ye) as nitrogen sources. The amount of EPS obtained also depended on the initial pH value of 4.5 and pH 7.0 (pH7.0, pH4.5). Based on the PCA, the optimum temperature for EPS production by the test strain was 20 °C (T_20), and the carbon source concentration was 30 g/L (Ccon_30). On the other hand, the obtained mycelial biomass (Biomass) was positively clustered with a temperature of 28°C (T_28) and carbon source concentrations of 15 g/L and 187.5 g/L (Ccon_15, Ccon_187.5). It is also worth noting that EPS synthesis did not correlate with biomass growth (Fig. 8A).

**Fig. 8.**
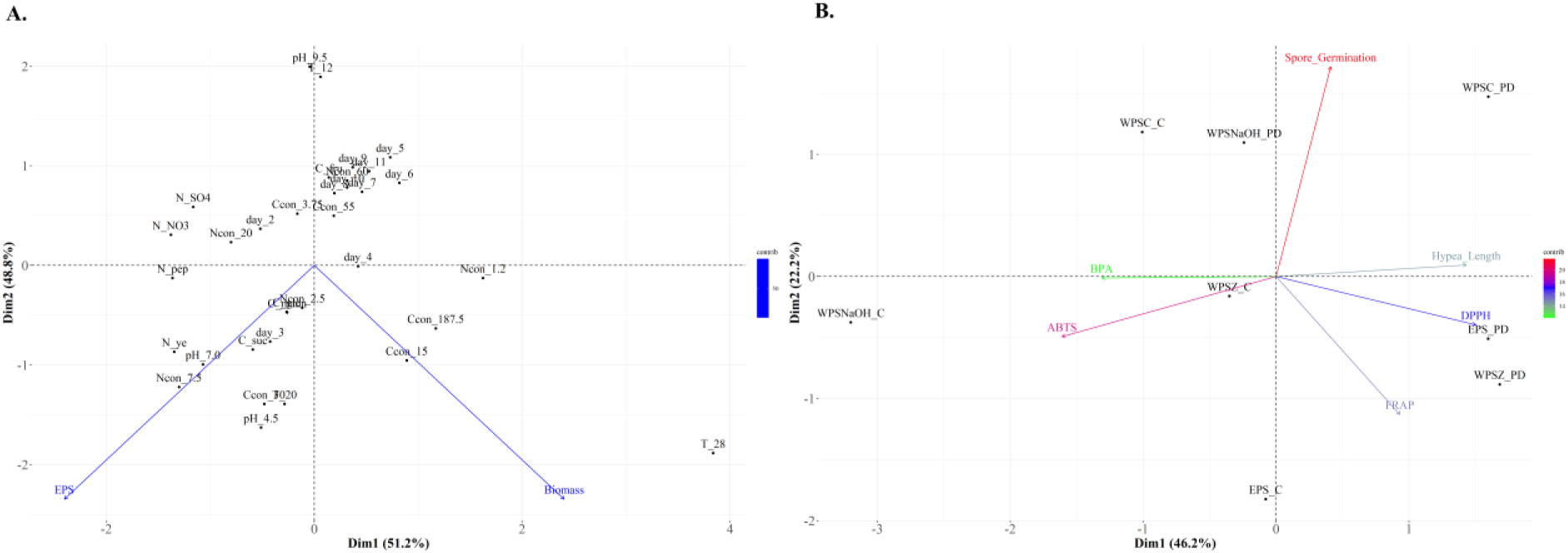
Biplot diagram of Principal Component Analysis (PCA) for: A. biomass and optimization parameters of EPS synthesis by *T. koningiopsis* strain growing on the medium with different carbon source: sucrose (C_suc), glucose (C_glc), fructose (C_fru), mannose (C_mann) and nitrogen source: peptone (N_pep), yeast extract (N_ye), NH4NO3 (N_NO3), and (NH4)2SO4 (N_SO4) with different concentration of carbon source (3.75, 15, 30, 55, 187.5 g/L) and nitrogen source (1.2, 2.5, 7.5, 20, 60 g/L) analyzed in ten days of incubation (2, 3, 4, 5, 6, 7, 8, 9, 10 and 11 days) at three temperatures (12, 20, 28 °C) and three pH value (4.5, 7.0, 9.5), B. The antioxidant properties (ABTS, DPPH, FRAP), bisphenol-A (BPA) binding capacity and antifungal properties (spore germination and hypea length of *F. culmorum* strain) of the PS frkations tested.

The relationship between the PS fractions obtained and their properties was assessed using Principal Component Analysis (PCA). The first two dimension of PCA explained 66.1% of the total variation, with principal component 1 (Dim1) accounting for 22.2% and principal component 2 (Dim2) accounting for 46,2% of the variance. ABTS antioxidant activity and BPA binding were positively correlated. These parameters were also strongly correlated with the PS fractions of WPSNaOH ***C***, WPSZ ***C***, and EPS ***C***. A similar correlation was observed between DPPH activity, FRAP, and the limiting shuttle length for the WPSZ ***PD*** and EPS ***PD*** fractions. In contrast, the spore germination of the *F. culmorum* strain did not correlate with the other parameters tested (Fig. 8B).

Student’s t-test was performed to analyse the relationship between antioxidant activities (ABTS, DPPH, and FRAP) and bisphenol-A (BPA) binding capacity between crude (***C***) and purified (***PD***) PS fractions (Table 4). For EPS activity, a significant difference was observed between the fractions (p<0.05), which shifted towards the ***C*** fraction. A similar relationship was observed for BPA binding, in which the ***C*** fractions also showed higher activity. In contrast, the ***PD***-purified fractions dominated DPPH activity, with only the WPSZ fraction showing no statistically significant difference between them. On the other hand, FRAP activity showed the greatest variability among the polymers tested. EPS and WPZ showed higher activity for the ***C*** fraction, and WPSC and WPSNaOH showed higher activity for the ***PD*** fraction.

**Table 4.** Two sample t-Student analysis of the relationship between crude (C) and purified (PD) PS for antioxidant and chelating properties.

| PS fraction | Properties |  |  |  |
| --- | --- | --- | --- | --- |
|  | ABTS | DPPH | FRAP | Chelating BPA |
| EPS (C) – EPS (PD) | t= 2.3373<br>p= **0.02545** | t= -3.5633<br>p= **0.001109** | t= 3.1675<br>p= **0.003241** | t= 1.346<br>p= 0.208 |
| WPSZ (C) – WSPZ (PD) | t= 2.3508<br>p= **0.02467** | t= -0.41742<br>p= 0.679 | t= -5.2484<br>p= **8.175e-06** | t= 8.2256<br>p= **9.222e-06** |
| WPSC (C) – WPSC (PD) | t= 3.9302<br>p= **0.0003953** | t= -2.7085<br>p= **0.0105** | t= -1.2968<br>p= 0.2034 | t= 2.4902<br>p= **0.03198** |
| WPSNaOH (C) – WPSNaOH (PD) | t= 3.8212<br>p= **0.0005389** | t= -3.6498<br>p= **0.0008722** | t= 0.86971<br>p= 0.3906 | t= 6.3675<br>p= **8.166e-05** |

## 4. Discussion

Strains belonging to the genus *Trichoderma* are extensively utilized as filamentous fungi in various applications. They are known for their contribution to agriculture and sustainable development by serving producers of numerous bioactive compounds, including EPS [16]. To achieve an adequate yield of EPS synthesis, optimisation of the medium and culture conditions is an important aspect of ongoing research. Literature data report that the most commonly used C sources are glucose and sucrose, and the most commonly used N sources are yeast extract or peptone. The optimum starting pH varies between 6.0-7.0 and the temperature is between 20-28°C. The greatest variability of the described strains was shown for the optimal culture time, where a discrepancy between the 4^th^ and 40^th^ days was observed [1,34]. In this study, we demonstrated that the *Trichoderma koningiopsis* strain is capable of synthesising EPS in liquid cultures. In the case of our isolate, the modified Czapek-Dox medium proved to be the most optimal, where the highest EPS synthesis efficiency was achieved with sucrose (30 g/L) and yeast extract (7.5 g/L) with a starting pH value of 7.0 and at 20°C. It seems that the described isolate showed optimum EPS synthesis after 3 days of culture. To date, it has been reported that strains of *T. pseudokoningii* [35], *T. harzianum* [20] and *T. viride* [36] are also capable of synthesising EPS. For *T. pseudokoningii* and *T. harzianum* strains, the optimal medium for synthesis was PDB medium, where the main source of C was glucose/starch, and the source of N was potato extract. The optimum temperature for these strains was 28°C, and they showed maximum synthesis after 10 d of incubation [20,35]. Interestingly, for the *T. viride* strain, the authors used a 25:75 mixture of Jensen’s and PDB medium, enriching the culture with sucrose (initial concentration 20 g/L) and mineral salts. However, the authors used stationary cultures at 28°C for 20 d [36]. Similar culture conditions were used for other Ascomycota strains. For *Fusarium culmorum*, *Leptosphaeria biglobosa*, and *L. maculans* strains, the modified Czapek-Dox medium also proved to be optimal. However, for these strains, peptone at a concentration of 7.5 g/L was the best source of N [25,37]. For Ascomycota strains belonging to the different types: *Eurotiomycetes Aspergillus* sp. DHE6 and *Sardoriomycetes* - *Cordyceps cicadae*, the optimal medium for EPS synthesis contained glucose (40 g/L and 2 g/L) and yeast extract (20 g/L and 0.3 g/L) at 28°C after 10 and 2 days, respectively [38,39]. On the other hand, for the *Mortierella alpina* strain (*Mucoromycota* type), the optimal C source was glucose (46.01 g/L), but the N source was already urea (7.48 g/L) at 25°C after 5 days of incubation [40]. The use of media with a high C and N content seems to be characteristic of Ascomycota strains. Moreover, the addition of organic N sources was more prevalent. In the case of our *T. koningiopsis* (*Hypocreales* order) strain, we obtained a synthesis yield of 1.17 g/L. The EPS yield classifies the described strain as a typical EPS producer [1,34]. A similar synthesis yield was achieved by strains representing different types of fungi: Ascomycota - the phytopathogenic strains *F. culmorum* (*Hypocreales* order) at 1.1 g/L and *Mucoromycota* - *M. alpina* at 1.51 g/L. Interestingly, the *F. culmorum* PGPF (Plant Growth Promoting Fungi) strain synthesised EPS at a level of 0.2 g/L and the *F. culmorum* DRMO (Deleterious Rhizosphere Microorganism) strain at a level of 0.7 g/L, which may suggest that EPS is involved in microorganism-plant interactions [25,40]. However, the variation in the efficiency of EPS synthesis by the Ascomycota strains was significantly greater. For example, the *Aspergillus* spp. DHE6, achieved an EPS synthesis yield of 26.1 g/L or *Aureobasidium pullulans* RYLF-10 with a synthesis yield of 40 g/L [39,41]. This indicates the need to search for new strains capable of synthesising bioactive EPS and optimising their culture. In addition to studies on EPS, studies on the properties of wall polymers (WPS) are of interest. Although the chemical composition of the fungal cell wall (FCW) is highly dependent on age, developmental stage and morphological state, many studies indicate a strong species dependence of the structure and proportions of individual wall components [42]. In our study, we obtained three fractions of WPS from the cultures: WPSZ (13.3%), WPSC (1.8%), and WPSNaOH (20.2%) using step-by-step extraction. However, the approaches for the extraction and study of wall polymers have varied. Jin and Ning (2013) isolated the fraction of water-soluble polymers by optimising the extraction time, ratio, and temperature. They were able to obtain a maximum of 10 514.7 µg/g of polysaccharides (PAF) from the cell wall of *Aspergillus fumigatus* AF1 strain, which is approximately 1.5 % the obtained mycelial weight, a value similar to that of our WPSC fraction. For the *Rhizoctonia solani* AG1 strain, the biomass was fractionated to obtain three PS fractions: the water fraction (WF), the cold alkaline fraction (CAF) and the hot alkaline fraction (HAF). The different fractions accounted for 5.66%, 8.05%, and 18.47% of the mycelial dry weight, respectively [44]. However, studies on the water-soluble wall polymer fraction of the *T. harzianum* strain indicated that the total amount of this fraction was more than 82.54% of the total mycelial weight [20]. On the other hand, chemical fractionation by alkaline (1 N NaOH) FCW extraction of three *F. culmorum* strains with different effects on plants (phytopathogenic, PGPF, DRMO) showed no statistically significant differences between the contents of the four fractions: (1) galacto-manno-glucan (alkali and water soluble), (2) glucan-chitin (alkali and water insoluble), (3) glucan weakly associated with chitin (alkali soluble at 20°C and water insoluble), (4) glucan strongly associated with chitin (alkali soluble at 70°C and water insoluble) [45].The aforementioned studies indicate that the variation in the amount of individual PS fractions within a single division of Ascomycota is highly variable but also dependent on the genus as well as on the species of the isolate under study.

The fractions obtained for both EPS and WPS differed not only in source but also in their structural properties. The purification process did not significantly affect the percentage composition of the monomers in the case of wall polymers. Glucose was predominant in all fractions. Only in the case of WPSC and WPSNaOH did the dominance of Glc increase in the ***PD*** friction. The greatest variability was observed in the case of EPS, where after purification, the amount of Glc in the EPS composition decreased by 10-fold and Man became the dominant monomer (Fig. 2). The EPS obtained from *T. harzianum* was also characterised by a high Man content, which accounted for 22.67% and 20.97% of the EPS1 and EPS2 fractions, respectively, but Glc was still dominant here. The four wall polymer fractions obtained by the authors exhibited variable Glc:Man ratios. In the higher molecular weight fractions (IPS1 and IPS2), Glc was dominant, in the IPS3 fraction, the ratio was close to 1:1, and in the lowest molecular weight fraction, Man was already starting to dominate. Ribose and arabinose were also present in both EPS and IPS (except for IPS3 and IPS4) [20]. In contrast, in *T. pseudokoningii*, galactose, glucose, mannose, xylose, rhamnose, and fucose were present in a molar ratio of 48.1:23.6:25.8:14.4:16.2:1 (Huang et al., 2012). GlcN and GalN were still observed in the EPS ***C*** of *T. koningiopsis* strain. In contrast, all PSs tested lacked clear arabinose or ribose signals, except for the WPSC fraction, where a small proportion of ribose and xylose was observed, as confirmed by FT-IR analysis. The composition of the cell wall fraction of *R. solani* also indicated a dominance of Glc, the content of which ranged from 73.35 to 96.16%, with the second dominant monomer already depending on the fraction described and being mannose, galactose, or fucose [44]. The sizes of the obtained fractions were also different between EPS and WPS. The EPS obtained had the largest fraction 52.48 kDa and the WPS in the range 158.49-831.76 kDa. EPS of strain *T. pseudokoningii*, where the EPS obtained was 31.9 kDa. However, the EPS obtained from cultured *Fusarium* strains had a higher molecular weight at 1.87*10^5^ Da for *F. solani* or 1000 kDa for *F. culmorum* strains [25,35,46]. The EPS FT-IR spectra contained bands characteristic of polysaccharides containing mostly mannose, galactose, and/or glucose monosaccharides. Previously, bands at 1416 and 1258 cm^-1^ were found for glucomannan and were assigned to the symmetrical stretching of COO groups whose vibration was modified by the presence of 4-O-methyl-α-D-glucuronic acid side groups [47]. While a band at 1050 cm^-1^ can be assigned to the ring vibrations overlapped by stretching vibration of (C-O-C) β-glycosidic. Bands at 884 and 807 cm^-1^ are also characteristic of glucomannans and can be assigned to the β-glycosidic linkage and stretching vibration of glucose, galactose, and mannose, respectively. The Raman spectra of EPS didn’t show many features but the main bands at 1458, 1380, 1335, 1260 cm^-1^, and bands in the range of 1200 – 1080 cm^-1^ can be resolved. The bands below 1200 cm^-1^ are characteristic for β-glycosidic linkage vibration for most of hemicellulosic polysaccharides. In the case of WPSZ and WPSC a similar band pattern in FT-IR spectra was obtained. The bands at 1148 and 993 cm^-1^ can be assigned to the xylan-type polysaccharides, and are attributed to the stretching and bending vibrations of C-O, C-C, and C-OH [48]. Moreover, the absorbance intensity in this region is highly influenced by the branching degree at O-2 and/or O-3 positions [49]. While bands at 1078 and 1015 cm^−1^ might be attributed to pyranose rings [50]. Previously, Černá et al. [51] assigned the specific wavenumbers to sucrose (995 cm^−1^), fructose (1064 and 1045 cm^−1^), arabinose (1052 and 979 cm^−1^), and mannose (1072 and 1033 cm^−1^). The bands with maxima in the range 1030–944 cm^−1^ were previously assigned to glucose. The spectral range of 1500 −1200 cm^-1^ is characteristic to C–H deformations, O–H bending, or the stretching of C–O bonds of carboxyl groups [52]. Bellow 950 cm^-1^ the information about the glycosidic linkages of polysaccharides can be found. The FT-IR band at 860 cm^-1^ was previously observed for the acetylated pectic polysaccharides but also for furanoid compounds in carbohydrates [53,54]. The Raman spectra of WPSZ and WPSC confirmed the presence of small amounts of proteins (bands between 1700-1500 cm^-1^). The bands at 1458, 1124, 1085, and 938 indicate presence of galactans [55]. On the other hand, band at 853 cm^-1^ is the most characteristic for α-type glycosidic linkage [56]. The bands at 1085 and 481 cm^-1^ were also found in acetylated glucomannan [55]. The FTIR spectra of WPSNaOH shows similar features to the other WPSes but the presence of bands at 1524 cm^-1^ and basically lack of significant bands below 900 cm^-1^ differ from those samples. While the maximum at 1030 cm^-1^ indicates presence of glucose and/or mannose [53]. The Raman spectra of WPSNaOH contain a band at 879 cm^-1^ which is characteristic for stretching of β-glycosidic linkage. Moreover, the bands at 1670 cm^-1^ indicates the presence of proteins (amid I). Linkages analysis indicated that the predominant bonds in the PSs tested were→4)-Hex II-(1→, likely in EPS and Glc in WPS. For EPS, WPSZ and WPSNaOH, the terminal monomers were Man and for WPSC it was Glc. A similar binding predominance was observed for polymers obtained from *T. harzianum* cultures, where →4)-α-_D_-Glc-(1→ residues were predominant [20]. In the *Craterellus cornucopioides* strain, the obtained EPSs was composed of terminal Man*p*-(1→ and branched →3,6)-Man*p*-(1→ residues, suggesting that these types of bonds are characteristic of polymers built from mannose subunits [57]. The wall polymers of *R. solani* were built mainly of →4)-Glc*p*-(1→ residues, however →6)-Glc*p*-(1→ residues were also present, which was also present in the WPSC fraction obtained from the cultured *T. koningiopsis* strain [44].

The polymers exhibited numerous bioactive properties with antioxidant as the most important ones.. PS obtained from cultures of the *T. koningiopsis* strain showed free radical scavenging ability in ABTS, DPPH, and FRAP assays. The activity of the obtained PSs increased with increasing concentration of the soluble sugar fraction. Interestingly, in some cases, the activity of the crude fraction (***C***) was higher than that of the purified fraction (***PD***). This may be related to the presence of phenolic compounds or low-molecular-weight fractions that were removed during purification. The WPSNaOH fraction had the highest ABTS free radical-scavenging ability, reaching 45%. For the DPPH radical, the EPS fraction had the highest scavenging capacity at 30%, and for FRAP, it was the WPSZ at the level of 6 µg/mL Fe^2+^. EPS obtained from *M. alpina* cultures at the same concentration (1 mg/mL) showed a similar level to that of the ABTS radical [40]. HUP-2 polymers obtained from *Hypsizygus ulmarius* cultures, on the other hand, achieved scavenging of 40% at a concentration of 1.5 mg/mL for the ABTS radical and at 2 mg/mL for the DPPH radical [58]. For the same radicals, the EPS1 fraction obtained from *T. harzianum* cultures showed a scavenging capacity of 50% at 1 mg/mL. In contrast, the other fractions oscillated at 10-20% radical scavenging capacity [20]. The reports described by the authors were similar to our results. Polymers obtained from the mycelia of *Auricularia* sp. also showed iron ion chelating capacity at similar levels[59]. Furthermore, the obtained PS fractions showed the ability to bind bisphenol-A in aqueous solutions. The WPSNaOH fractions proved to be the most efficient, binding up to 75% of BPA. The removal of BPA from the environment is one of the most important challenges, owing to its high toxicity and negative environmental impact. Studies available in the literature indicate that microbially derived EPS are involved in the binding of BPA and can be used in the removal of this compound from water, for example, in wastewater treatment plants [60]. The most intriguing property of the obtained PSs was their ability to limit the growth of the phytopathogenic strain *F. culmorum*. The obtained fractions either interacted directly with spore germination or limited the length of the germinated hyphae. The EPS and WPSZ fractions were the most effective at limiting germination. In contrast, the WPSC and WPSNaOH fractions were the most effective in limiting the length of the germinated hyphae. Similar properties were exhibited by polysaccharides obtained from *Oudemansiella radicata*, where, already at a concentration of 0.5 g/L, the authors observed inhibition of the growth of the *Penicillium digitatum* strain infesting oyster mushroom (*Pleurotus ostreatus*) during storage. The authors indicated that the OWRPs obtained affected the cell wall of the fungus, causing cell deformation and cell death [61]. Similar properties against the *F. oxysporum* strain were shown by EPS obtained from *Porphyridium sordidum* cultures, where inhibition of phytopathogen growth on *Arabidopsis thaliana* leaves was observed after the application of a concentration of 2 mg/mL. Furthermore, the authors indicated that EPS could stimulate response markers in plant tissues [62].

## 5. Conclusions

The *Trichoderma koningiopsis* strain has the capability to produce extracellular polymers (EPS), a feature that holds significance concerning the potential applications of this species in biopreparations and its utilization in diverse agricultural practices.. As part of the ongoing culture optimisation, we were able to increase the synthesis yield to 1.17 g/L. In addition, three fractions of WPS wall polymers were obtained in our study: the cold water soluble (WPSZ), hot water soluble (WPSC) and alkali soluble (WPSNaOH) fractions, which accounted for 13.3%, 1.8% and 20.2% of the mycelial dry weight, respectively. The EPS obtained was composed mainly of mannose, and the WPS were glucans. PS fractions were predominantly built of →4)-Glc- (1→ linked residues, with branching at →3,6)- as well as →4,6)-positions. FT-IR and FT-Raman analyses showed that α-bonds dominate in the WPSZ and WPSC fractions, whereas β-bonds dominate in the EPS and WPSNaOH fractions. They showed the ability to scavenge free radicals using the ABTS, DPPH, and FRAP methods, indicating their potential for use as antioxidant compounds in medicine and the food industry. Moreover, their ability to adsorb BPA suggests their utility in water purification systems. However, according to us, the most important factor is their ability to reduce the growth of the phytopathogenic strain *F. culmorum*, which is one of the most common causes of agricultural loss. In addition, it shows the ability to synthesise mycotoxins; therefore, it is important to limit their appearance in crops. *T. koningiopsis* has the ability to limit the growth of *Fusarium* strains in direct interactions. By proving that it can also inhibit the growth of phytopathogens through indirect interactions, this opens new perspectives for the design and manufacture of biopreparations.

## CRediT authorship contribution statement

**Artur Nowak**: Writing – review & editing, Writing – original draft, Visualization, Methodology, Investigation, Data curation, Formal analysis, Conceptualization, Funding acquisition, Project administration, **Kamila Wlizło**: Investigation, Methodology **Iwona Komaniecka**: Writing – review & editing, Writing – original draft, Methodology, Investigation, **Monika Szymańska-Chargot**: Writing – original draft, Methodology, Investigation, Visualization, **Artur Zdunek**: Writing – review & editing, Methodology, **Justyna Kapral-Piotrowska**: Methodology, Investigation, **Katarzyna Tyśkiewicz**: Writing – review & editing, **Jolanta Jaroszuk-Ściseł**: Supervision, Writing – review & editing, Methodology.

## Declaration of competing interest

The authors declare that they have no known competing financial interests or personal relationships that could have appeared to influence the work reported in this paper.

## Data availability

All data are available in the Zenodo repository: https://doi.org/10.5281/zenodo.12785609

## Acknowledgements

GA was created with BioRender.com

## Fundings

The research was funded by National Science Center Poland: Miniatura 6 (2022/06/X/NZ9/00569)

## Appendix A.

Supplementary data

## Notes

### Competing Interest Statement

The authors have declared no competing interest.

