## supplementary table for "Isolation and characterization of cell wall and extracellular polysaccharides from cultures of the mycoparasitic strain *Tirochoderma koningiopsis*"

**Title**

**Authors and affiliations**

Artur Nowak^1*^, Kamila Wlizło^1^, Iwona Komaniecka^2^, Monika Szymańska-Chargot^3^, Artur Zdunek^3^, Justyna Kapral-Piotrowska^4^, Katarzyna Tyśkiewicz^5^, Jolanta Jaroszuk-Ściseł^1^

^1^Department of Industrial and Environmental Microbiology, Institute of Biological Sciences, Maria Curie-Skłodowska University, Akademicka 19, 20-033 Lublin, Poland;

^2^Department of Genetics and Microbiology, Institute of Biological Sciences, Maria Curie-Skłodowska University, Akademicka 19, 20-033 Lublin, Poland;

^3^Institute of Agrophysics, Polish Academy of Sciences, Doświadczalna 4, 20-290 Lublin, Poland;

^4^Department of Functional Anatomy and Cytobiology, Institute of Biological Sciences, Maria Curie-Sklodowska University, Akademicka 19, 20-033 Lublin, Poland;

^5^Łukasiewicz Research Network- New Chemical Syntheses Institute, Al. Tysiąclecia Państwa Polskiego 13A, Puławy 24-110, Poland,

*** Corresponding author**

^*^Artur Nowak

^1^Department of Industrial and Environmental Microbiology, Institute of Biological Sciences, Maria Curie-Skłodowska University

Akademicka 19, 20-033 Lublin, Poland

Table S1. Results of step-by-step optimisation of T. koningiopisis strain culture on the biomass obtained (g/L). Statistical analysis was performed by Anova test using Tukey post hoc p<0.05.

|  | **Optimised parameter** | | | | | | | | | | | | | | | | | |
| --- | --- | --- | --- | --- | --- | --- | --- | --- | --- | --- | --- | --- | --- | --- | --- | --- | --- | --- |
| **Biomass**  **g/L** | **Day of culture** | | | | | | | | | | | | | | | | | |
|  | 2 | 3 | 4 | | | 5 | | 6 | | 7 | 8 | | | 9 | | 10 | | 11 |
|  | 10.26±0.38  d | 15.37±0.73  a | 15.75±0.78  a | | | 12.48±0.65  bcd | | 13.91±0.81  ab | | 12.79±0.37  bc | 11.74±0.01  bcd | | | 11.39±1.04  cd | | 12.02±0.71  bcd | | 12.21±1.46  bcd |
|  | **Carbon source** | | | | | | | | | | | | | | | | | |
|  | Sucrose | | | | Glucose | | | | | Fructose | | | | | Mannose | | | |
|  | 15.01±0.07  a | | | | 14.79±0.13  a | | | | | 10.86±0.08  b | | | | | 14.84±0.07  a | | | |
|  | **Nitrogen source** | | | | | | | | | | | | | | | | | |
|  | Peptone | | | | Yeast extract | | | | | NH_4_NO_3_ | | | | | (NH_4_)_2_SO_4_ | | | |
|  | 8.8±0.31  b | | | | 11.95±0.11  a | | | | | 6.93±0.41  c | | | | | 6.65±0.12  c | | | |
|  | **Temperature (°C)** | | | | | | | | | | | | | | | | | |
|  | 12 | | | | | | 20 | | | | | | 28 | | | | | |
|  | 6.32±0.79  c | | | | | | 18.55±3.26  b | | | | | | 37.78±3.65  a | | | | | |
|  | **Initial pH value** | | | | | | | | | | | | | | | | | |
|  | 4.5 | | | | | | 7.0 | | | | | | 9.5 | | | | | |
|  | 18.59±0.15  a | | | | | | 13.63±4.56  a | | | | | | 5.46±1.12  b | | | | | |
|  | **Carbon source concentrations (g/L)** | | | | | | | | | | | | | | | | | |
|  | 3.75 | | | 15 | | | | | 30 | | | 55 | | | | | 187.5 | |
|  | 11.11±0.49  c | | | 21.64±0.92  a | | | | | 17.74±0.94  b | | | 12.66±0.58  c | | | | | 21.46±0.42  a | |
|  | **Nitrogen source concentrations (g/L)** | | | | | | | | | | | | | | | | | |
|  | 1.2 | | | 2.5 | | | | | 7.5 | | | 20 | | | | | 60 | |
|  | 21.24±0.96  a | | | 15.24±0.96  b | | | | | 13.62±0.7  bc | | | 9.64±0.64  d | | | | | 11.73±.031  c | |
